# Defining proximity proteomics of post-translationally modified proteins by antibody-mediated protein A-APEX2 labeling

**DOI:** 10.1101/2021.03.05.434178

**Authors:** Xinran Li, Jiaqi Zhou, Wenjuan Zhao, Qing Wen, Weijie Wang, Huipai Peng, Kelly J. Bouchonville, Steven M. Offer, Zhiquan Wang, Nan Li, Haiyun Gan

## Abstract

Proximity labeling catalyzed by promiscuous enzymes, such as APEX2, has emerged as a powerful approach to characterize multiprotein complexes and protein-protein interactions. However, current methods depend on the expression of exogenous fusion proteins and cannot be applied to post-translational modifications. To address this limitation, we developed a new method to label proximal proteins of interest by antibody-mediated protein A-APEX2 labeling (AMAPEX). In this method, a modified protein is bound *in situ* by a specific antibody, which then tethers a protein A-APEX2 (pA-APEX2) fusion protein. Activation of APEX2 labels the nearby proteins with biotin; these proteins are then purified using streptavidin beads and are identified by mass spectrometry. We demonstrate the utility of this approach by profiling the binding proteins of histone modifications including H3K27me3, H3K9me3, H3K4me3, H4K5ac and H4K12ac, and we verified the genome-wide colocalization of these identified proteins with bait proteins by published ChIP-seq analysis. Overall, AMAPEX is an efficient tool to identify proteins that interact with modified proteins.

## Introduction

Biological functions are regulated by interacting biomolecules (protein, DNA, RNA, etc.); dysregulation of these interactions can lead to human diseases including cancers^1,2^. Methods that map these molecular interactions provide tools to study biological processes and therapeutics for human diseases. Recently, proximity labeling was developed and used to map molecular interactions. Proximity labeling uses engineered enzymes, such as peroxidase or biotin ligase, that are tagged to a protein of interest^3^ to modify nearby factors that interact with that protein of interest. This method has been successfully utilized to map protein-protein^3^, RNA-protein^4–6^, protein-DNA^7,8^, and chromatin interactions^9^. However, proximity labeling has some limitations. First, the expression of exogenous fusion proteins with engineered enzymes is required, which limits its use in difficult-to-transfect cells and tissues. Additionally, mapping the biomolecules that interact with posttranslationally modified (PTM) proteins like histones is complex.

Histone modifications play critical roles in regulating basic biological processes to maintain cell identity and genome integrity^10,11^. These modifications are recognized by reader proteins and then form functional multiprotein complexes with other regulatory factors in a spatiotemporal manner^12^. Alterations in the interacting networks of the histone modifications can lead to human diseases^13^. However, current proteomics-based assays to measure the affinity of proteins to chromatin marks^14–18^ rely on the use of synthetic histone peptides, *in-vitro*-reconstituted nucleosomes, or expression of external protein domains; thus, it is challenging to identify the proximal protein interactome of histone modifications *in situ*.

Here, we overcome the limitations of traditional proximity labeling methods using a protein A-APEX2 (pA-APEX2) fusion protein. The protein of interest is bound *in situ* by a specific antibody, which then tethers a pA-APEX2 fusion protein. Activation of APEX2 labels the nearby proteins with biotin; these proteins are then purified using streptavidin beads and identified using mass spectrometry.

## Results

The strategy behind antibody-mediated pA-APEX2 labeling (AMAPEX) is to tether a peroxidase or biotin ligase to antibodies that are specifically bound to a protein of interest (here, histone modifications; Fig. 1a). Subsequent activation of the tethered enzyme should result in biotinylation of biomolecules near the target protein. Identification of the biotinylated proteins by mass spectrometry is expected to provide information about the nearby proteomic landscape of the protein of interest (Fig. 1b). We selected APEX2 as the enzyme of choice because it has robust enzymatic activity *in vitro* and can be stringently controlled by H_2_O_2_^5^.

**Figure 1.**
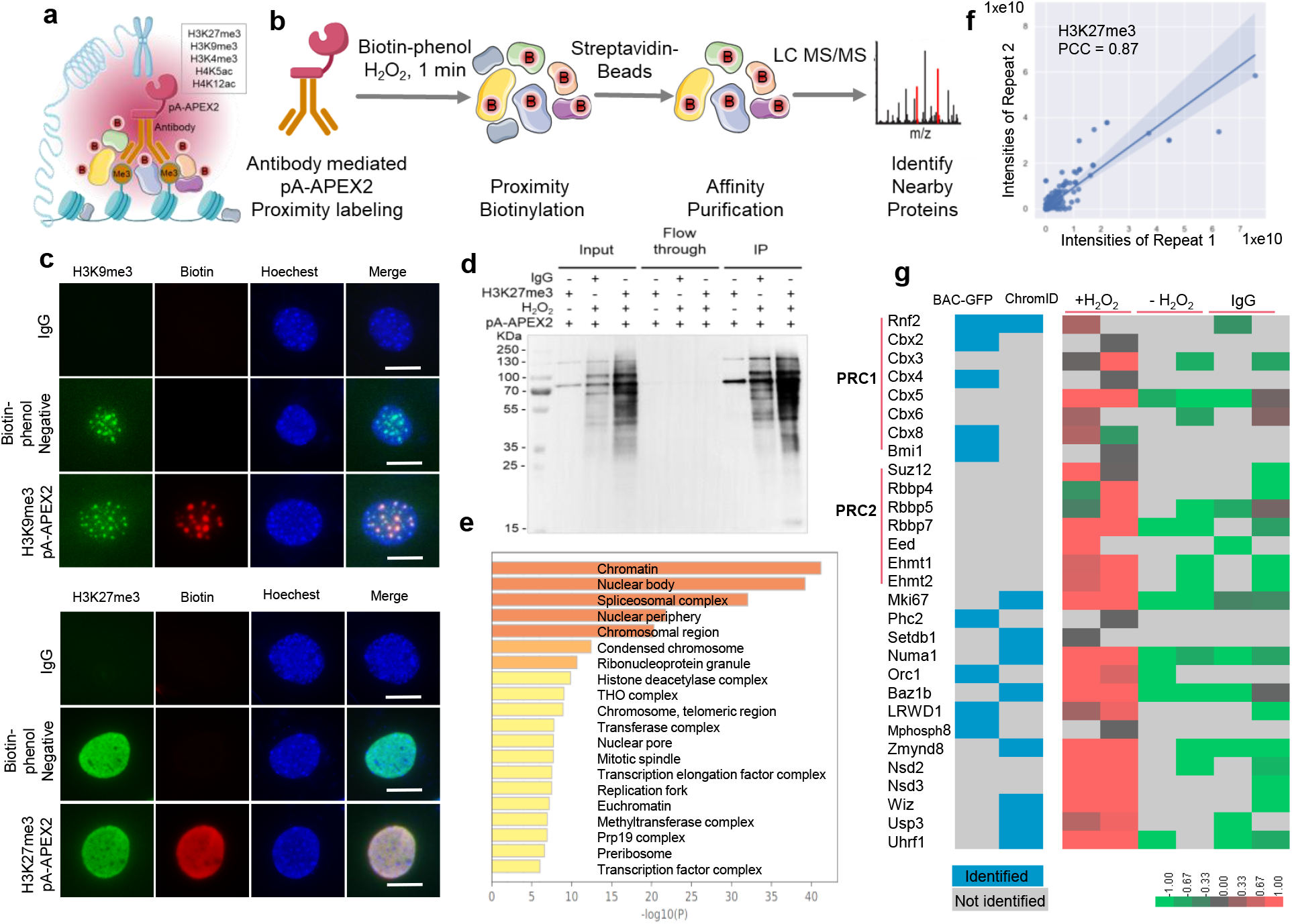
Antibody-mediated proximity biotinylation by pA-APEX2. **a**, Illustration of antibody-mediated pA-APEX2 proximity labelling. pA-APEX2 is recruited to the targeting sites by specific histone modification antibodies. Biotin-phenol and H_2_O_2_ are added to 0.1% formaldehyde-fixed cells for 1 minute to induce biotinylation (B, biotin) of proteins within 20 nm of APEX2. **b**, Biotinylated proteins are purified using streptavidin beads and analyzed by LC MS/MS. **c**, Fluorescence imaging of histone modifications and antibody-mediated biotinylation. H3K9me3 and H3K27me3 were visualized by immunofluorescence staining. Biotinylation was induced by adding biotin-phenol and H_2_O_2_ and visualized by staining with streptavidin-Cy3 (red). Nuclei were counterstained with Hoechest33342. Scale bars, 10 μm. **d**, pA-APEX2-mediated protein labeling in whole-cell lysates. Whole-cell extracts from MEF cells were incubated with pA-APEX2 and H3K27me3 antibody, and biotinylation was induced by adding biotin-phenol and H_2_O_2_ and analyzed by western blot as indicated. **e**, The top 20 enriched cellular components GO terms of H3K27me3-proximal proteins. Bar plots represent the -log10(p value) of enriched terms. **f**, The reproducibility of two biological replicates of pA-APEX2 experiments in the identification of H3K27me3-interacting proteins was calculated using the Pearson correlation coefficient (PCC). **g**, H3K27me3-interacting proteins identified by pA-APEX2. Heatmap showing the enrichment as log2-fold intensity change of interacting proteins relative to the controls as indicated. Data are shown as Z scores. Blocks in blue represent the enrichment of proteins identified by ChromID and BAC-GFP in previous publications.

We confirmed the enzymatic activity of the purified pA-APEX2 by labeling the whole-cell lysate *in vitro.* (Supplementary Fig. 1a-d). We then modified the immunofluorescence assay to test if enzymatically active pA-APEX2 can be recruited to the protein of interest by specific antibodies. The permeabilized cells were incubated with H3K9me3/H3K27me3 antibodies followed by pA-APEX2. The unbound pA-APEX2 was extensively washed out, and biotinylation was induced by H_2_O_2_ and biotin-phenol (BP). The co-localization of biotin and H3K9me3/H3K27me3 suggested that pA-APEX2 can be recruited to specific sites by specific antibodies and activated *in situ* (Fig 1c). As expected, no enrichment was observed in the samples without BP or in the IgG controls. We then tested the activity of pA-APEX2 in cell suspensions. Lightly crosslinked MEF cells were permeabilized and incubated with H3K27me3 antibody followed by pA-APEX2 (see detailed description in the Methods section). Biotinylation of the proteins around H3K27me3 was induced and then analyzed by western blot (Supplementary Fig.1e). Efficient biotin labeling in the presence of H3K27me3 antibody indicated that pA-APEX2 targeting can be accurately controlled. To test whether pA-APEX2 could be used to identify proteins associated with histone modifications *in situ,* the biotinylated proteins were enriched with streptavidin beads and analyzed using quantitative LC-MS/MS (Fig.1 b and d); samples that were incubated with IgG and without H_2_O_2_ were included as negative controls. Compared to ChromID^14^ and BAC-GFP^15^, we reproducibly identified most of the PRC1 and PRC2 subunits using quantitative LC-MS/MS (Fig. 1f, g). In addition to the known proteins, we identified a number of candidate proteins associated with H3K27me3 (Supplementary Table 1). Gene ontology (GO) term analysis indicated that these proteins were enriched in known cellular components associated with H3K27me3, including transcription repressor complex^19^, histone methyltransferase complex^19^, and DNA replication fork^20,21^ (Fig. 1e and Supplementary Fig. 2). To validate our results, we assessed the genomewide enrichment of three candidates, the H3K36 methyltransferase Nsd2 and the polybromo-associated BAF (PBAF) subunits Brd7 and Arid2^22^, by analyzing published ChIP-seq datasets^23^ (Supplementary Fig. 3). We verified the colocalization of the PBAF subunits Brd7 and Arid2 with H3K27me3 in MCF-7 cells by ChIP-seq (Supplementary Fig 3a-d), which indicates that PBAF may recognize H3K27me3 to remodel suppressed chromatin^24^. The ChIP-seq results also showed that around half of Nsd2 peaks in K-562 cells overlap with H3K27me3 peaks (Supplementary Fig 3e, f), which supported our results that Nsd2 is proximal to H3K27me3. H3K36me2 is a negative regulator of H3K27me3^25^ and has been boundary barrier of PRC-mediated H3K27me3 spreading^26^. Thus, our results suggest that the binding of NSD2 to H3K27me3 may provide new insights into the mechanism that regulates epigenetic spreading.

We next generalized our method to map the interactomes of major histone modifications including H3K4me3, H3K9me3, H4K5ac, and H4K12ac. Western blot demonstrated successful labelling and enrichment of the proximal proteins of these histone marks (Fig. 2a and Supplementary Fig. 4). We next performed LC-MS/MS to identify and map the proteomes of these modifications. We first analyzed the proteins enriched by H3K9me3 labeling and identified several known H3K9me3 binding proteins including the reader proteins Lrwd1^27,28^, Cbx5 (HP1α) and Cbx3 (HP1γ)^29^; the H3K9 methyltransferases Ehmt1 and Setdb1; the DNA replication–dependent nucleosome assembly chaperones Chaf1a and Chaf1b; and the DNA methylation maintenance proteins Mecp2, Uhrf1, and DNMT1 (Supplementary Fig. 5a, b and Supplementary Table 1). We also identified H3K9me3-binding proteins that were enriched in functional complexes like the DNA replication fork, methyltransferase complex, and PcG protein complex^30^ (Supplementary Fig 5c, d). Therefore, this method could help us understand the mechanism of epigenetic inheritance including histone and DNA methylation.

**Figure 2.**
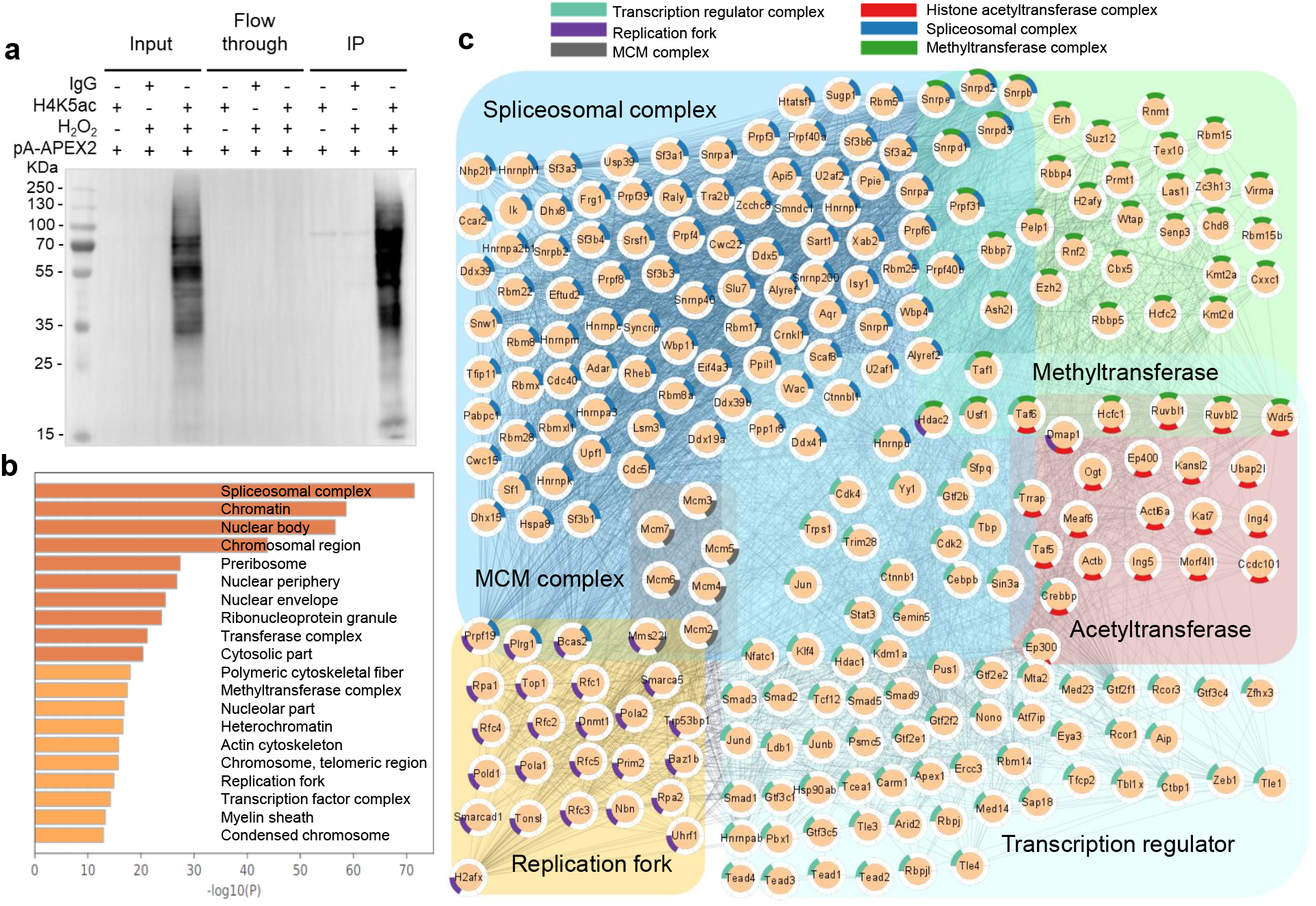
The proximal proteome of H4K5ac identified by AMAPEX. **a**, H4K5ac-proximal proteins were biotinylated by AMAPEX and purified using streptavidin beads. Whole-cell lysates, flow through, and streptavidin-purified proteins were analyzed by western blot as indicated. **b**, The top 20 enriched cellular component GO terms for the H4K5ac-proximal proteins. Bar plots represent the -log10(p value) of the enriched terms. **c**, Network analysis of H4K5ac interactomes according to the major cellular component GO terms (N proteins = 254, of all 1783). Individual proteins are shown as nodes and interactions are shown as edges. The interactions were retrieved from the STRING database with interaction score > 0.4. Proteins were selected based on a minimum of log2-FC = 1 in two pA-APEX2 experiments.

In addition to the above, we also identified the MLL H3K4 methyltransferase Kmt2a and COMPASS-related proteins including Wdr5 and Cxxc1^31^ in the H3K4me3 interactome (Supplementary Fig. 6a, b and Supplementary Table 1). H3K4me3-proximal proteins are mostly enriched in pathways related to active transcription including RNA splicing and euchromatin (Supplementary Fig. 6c, d).

We next focused on proteins that interacted with H4K5ac, the histone modification that marks newly synthesized histones^32^. The H4K5ac interactome is enriched with MCM complex^33^ and DNA replication fork proteins (Fig. 2b, c), which confirms that newly synthesized H3/H4 complexes are deposited in a DNA replication–dependent manner. However, we did not observe an association between H4K12ac-labeled proteins and DNA replication (Supplementary Fig. 7a-c), which suggested that the H4K12ac modification may not be recognized by new histone H3/H4 deposition machinery. Both H4K5ac and H4K12ac are enriched in RNA spliceosomes (Fig. 2b, c and Supplementary Fig. 7a, b), indicating that they may be involved in RNA splicing.

In summary, we provide a new approach to identify proximal complexes of a protein of interest without the expression of exogenous fusion proteins. We applied our method to identify proteins associated with major histone modifications (Supplementary Fig. 8 and Supplementary Table 2). Both known and previously unreported interactors of these histone modifications were identified by our method. APEX2 can also biotinylate RNA and DNA^34^, which provides the opportunity to apply AMAPEX to map the DNA and RNA molecules that bind to a protein of interest in future studies.

## Materials

| Antibodies | SOURCE | IDENTIFIER |
| --- | --- | --- |
| H3K4me3 | Active Motif | Cat# 39159; RRID: AB_2615077 |
| H3K9me3 | Abcam | Cat# ab8898; RRID: AB_306848 |
| H3K27me3 | Cell Signaling<br>Technology | Cat# 9733S; RRID: AB_2616029 |
| H4K5ac | Abcam | Cat# ab51997; RRID: AB_2264109 |
| H4K12ac | Abcam | Cat# ab177793; RRID: AB_2651187 |
| IgG | Abcam | Cat# ab6701; RRID:AB_956011 |
| Donkey anti-Rabbit IgG<br>(H+L) Alexa Fluor Plus<br>488 | Invitrogen | Cat# A32790; RRID: AB_2762833 |
| Streptavidin-Cy3 | Sigma | Cat# S6402-1ML |

## Methods

### Plasmids

The 3XFlag-pA-Tn5-Fl plasmid (Addgene plasmid # 124601) was used as the backbone to construct 3XFlag-pA-APEX2. Tn5 cDNA was cleaved with *NdeI* and *Spe*I and replaced with APEX2, which was amplified by polymerase chain reaction (PCR) from GFP-APEX2-NIK3x (GFP-APEX2-NIK3x was a gift from Alice Ting (Addgene plasmid # 129274; http://n2t.net/addgene:129274; RRID: Addgene_129274).

### 3XFlag-pA-APEX2 protein purification

Protein purification was performed as described by Steven Henikoff *et al.^35^*. The 3XFlag-pA-APEX2 plasmid was transformed into C3013 cells and incubated overnight at 37°C. A single colony was selected and inoculated into 3 mL LB medium, and growth was continued at 37°C for 4h. This culture was used to start a 400-mL culture in 100μg/mL carbenicillin-containing LB medium and incubated on a shaker until it reached O.D. ~0.6; the culture was then chilled on ice for 30 min. Fresh IPTG (Sigma, I6758) was added to 0.25 mM to induce expression, and the culture was incubated at 18°C on a shaker overnight. The culture was collected by centrifugation at 6,000 × *g* and 4°C for 30 min. The pellet was stored at −80°C until processing. The protein purification steps were described as follows. Briefly, a frozen pellet was resuspended in 40 mL chilled HEGX Buffer (20 mM HEPES-KOH at pH 7.2, 1 M NaCl, 1 mM EDTA, 10% glycerol, 0.2% Triton X-100) including 1× Roche Complete EDTA-free protease inhibitor tablets (Invitrogen, M9260G) and kept on ice for 15 min. The lysate was sonicated 15 min (300 W, 3 s on, 5 s off) on ice. The sonicated lysate was centrifuged at 16,000 × *g* at 4°C for 30 min, and the soluble fraction was moved to fresh 50-mL tubes. A 4-mL aliquot of chitin resin (NEB, S6651S) was packed into each of two disposable columns (Bio-rad, 7321010). Columns were washed with 20 mL HEGX Buffer. The supernatant was added to the chitin resin slowly and then incubated on a rotator at 4°C for 1 h. The unbound soluble fraction was drained, and the columns were washed with 20 mL HEGX buffer and with 20 mL HEGX buffer containing Roche Complete EDTA-free protease inhibitor tablets. The chitin slurry was transferred to a 15-mL tube and resuspended in elution buffer (6 mL HEGX buffer with 100 mM DTT (Roche, D0632)). The tube was placed on a nutator at 4°C for 60 h. The eluate was collected and dialyzed twice in 1 L Dialysis Buffer (100 mM HEPES-KOH pH 7.2, 0.2 M NaCl, 0.2 mM EDTA, 2 mM DTT, 0.2% Triton X-100, 20% Glycerol). The dialyzed protein solution was concentrated using Amicon Ultra-4 Centrifugal Filter Units 30 K (Millipore UFC803024), and sterile glycerol was added to make a final 50% glycerol stock of the purified protein. The purified protein was aliquoted and stored at −20°C. The pA-APEX2 purification was analyzed by SDS-PAGE. The concentration of pA-APEX2 was determined using BSA standards.

### Mammalian cell culture

Mouse embryonic fibroblast (MEF) cell lines were cultured in DMEM/high glucose (HyClone, SH30243.01) supplemented with 10% fetal bovine serum (ExCell Bio, FSP500), 100 units/mL penicillin, and 100 mg/mL streptomycin at 37°C under 5% CO_2_. Mycoplasma testing was performed before experiments.

### Immunofluorescence staining and fluorescence microscopy

MEF cell lines were fixed with 4% paraformaldehyde in PBS at room temperature for 15 min. Cells were then washed with PBS three times and blocked for 1 h with 3% BSA in 0.1% PBST (blocking buffer) at room temperature. Cells were incubated with primary antibodies (Rabbit anti-H3K9me3 antibody, Abcam, ab8898, RRID: AB_306848, 1:100 dilution; Rabbit anti-H3K27me3 antibody, Cell Signaling Technology, 9733S, RRID: AB_2616029, 1:100 dilution) in blocking buffer for 1 h at room temperature. After washing three times with PBS, cells were incubated with pA-APEX2 (in this study, 4 μg/μL, 1:400) in blocking buffer for 1 h and then washed three times with PBST. Next, cells were incubated with 500 μM biotin phenol (BP) in PBS at room temperature for 30 min. H_2_O_2_ then was added to each well to a final concentration of 1 mM, and the plate was gently agitated for 1 min. The reaction was quenched with an equal volume of 2 × quench buffer (10 mM Trolox, 20 mM sodium ascorbate, and 20mM sodium azide in PBS). Samples incubated with IgG and without BP (no-BP) were included as negative controls. After washing three times with PBS, cells were incubated with secondary antibodies (Alexa Fluor 488 (Invitrogen, A32790, 1:200 dilution) and streptavidin-Cy3 (Sigma, S6402-1ML, 1:300 dilution)) in blocking buffer for 1 h at room temperature. Cells were washed and incubated Hoechst for 10 min at room temperature, washed three times with PBS, and imaged.

### pA-APEX labeling *in vitro*

Total protein of MEF cell lines was extracted with RIPA lysis buffer (50 mM Tris-HCl pH 7.5, 150 mM NaCl, 1.5 mM MgCl_2_, 1 mM EGTA, 0.1% SDS, 1% NP-40, 0.4% sodium deoxycholate, 1 mM DTT, 1mM PMSF, and 1× Roche Complete EDTA-free protease inhibitor tablets) for 15 min at 4°C. Cell extracts were sonicated (100 W, 3 s on, 3 s off) for 3 min. Cell extracts were clarified by centrifugation, and the amount of protein in each supernatant was measured. 20 μg total protein was incubated with 10 μM pA-APEX2 and 0.5 mM BP in PBS for 1 min. The reaction was triggered by mixing with 1 mM H_2_O_2_ and stopped with quench buffer. No-H_2_O_2_, no-BP, and no-pA-APEX2 samples were included as negative controls.

### pA-APEX labeling in MEF cell lines

The labeling is adapted and modified from the Cut&tag method^36^. 2×10^7^ cells were washed with 10 mL PBS, pooled into a 50-mL tube, and centrifuged at 250 × *g* for 5 min. The cell pellet was resuspended in 1 mL PBS, crosslinked with freshly prepared formaldehyde at a final concentration of 0.1% at room temperature for 15 min, and quenched with 1/10 volume of 1.25 M glycine. The tube was inverted several times, shaken gently for 5 min, and centrifuged at 500 × *g* for 5 min. The pellet was then washed once with 10 mL PBS and centrifuged at 500 × *g* for 5 min. Supernatant was carefully aspirated, and the cell pellet was resuspended in 1 mL wash buffer (20 mM HEPES pH 7.5, 150 mM NaCl, 0.5 mM spermidine, 1× Protease Inhibitor EDTA-Free tablet (Invitrogen, M9260G)), transferred to a 1.5-mL tube, and centrifuged at 500 × *g* for 5 min. The pellet was resuspended in 300 μL antibody buffer (4 μL 0.5 M EDTA, 3.3 μL 30% BSA, and 10 μL 5% digitonin in 1 mL wash buffer) with 2 μL primary antibody (H3K27me3, H3K4me3, H3K9me3, H4K5ac and H4K12ac) and incubated overnight at 4°C. Then, the pellets were washed twice with 1 mL 0.01% digitonin wash buffer (20 μL 5% digitonin in 10 mL wash buffer) and centrifuged at 500 × *g* for 5 min. The pellet was incubated with 300 μL 500 μM BP in digitonin wash buffer for 30 min before incubation with 3 μL 100 mM H_2_O_2_ in wash buffer for 1 min (final concentration of 1 mM H_2_O_2_). The reaction was quenched by adding 300 μL 2× quench buffer (20 mM sodium azide, 20 mM sodium ascorbate, 10 mM Trolox in wash buffer)^37^. Then, the pellet was washed twice with quench buffer. After carefully aspirating the supernatant, the cell pellet was flash frozen and stored at −80°C until use.

### Streptavidin pull-down of biotinylated proteins and western blot analysis

pA-APEX2-labeled cell pellets were lysed in RIPA lysis buffer (50 mM Tris-HCl pH 7.5, 150 mM NaCl, 1.5 mM MgCl_2_, 1 mM EGTA, 1% SDS, 1% NP-40, 0.4% sodium deoxycholate, 1 mM DTT, 1mM PMSF, and 1× Roche Complete EDTA-free protease inhibitor tablets) for 15 min on ice. Cell extracts were sonicated (100 W, 3 s on, 3 s off) for 3 min and then boiled for 10 min at 100°C. Cell extracts were clarified by centrifugation, and the amount of protein in each supernatant was measured. 5% of the supernatant was saved as input for western blot analysis. SDS in the sample was diluted to 0.2% with 1x cold RIPA buffer (RIPA buffer without SDS). Streptavidin–Sepharose beads (GE Healthcare, 17511301) were washed twice with 1x cold RIPA buffer (0.2% SDS), and 800 μg of each sample was separately incubated with 50 μL bead slurry with rotation for 4 h at 4°C. 5% of the flow through was saved for western blot analysis. The beads were subsequently washed twice with 1 mL wash buffer (50 mM Tris-HCl pH 7.5, 1% SDS), twice with 1 mL RIPA wash buffer (50 mM Tris-HCl pH 7.5, 150 mM NaCl, 1.5 mM MgCl_2_, 1 mM EGTA, 0.2% SDS, 1% NP-40, 1 mM DTT), twice with 1 mL 8 M urea buffer, twice with 1 mL 30% acetonitrile, and twice with 1 mL 20 mM ammonium bicarbonate. 5% of the beads were saved for western blot analysis, and the remaining beads were used for LC-MS/MS analysis. For western blot analysis, biotinylated proteins were eluted by boiling the beads in 10 μL 5 × protein loading buffer and separated by 10% SDS-PAGE. The proteins were transferred to 0.22 μm PVDF membrane (Millipore) and stained with Ponceau S. The blots were then blocked in 1% BSA in TBST at room temperature for 1 h and stained with streptavidin-HRP (Beyotime, A0303, 1:5000 dilution) in TBST for 1 h at room temperature. Blots were then washed with TBST buffer three times for 5 min, developed with Clarity Western ECL substrate (Thermofisher), and imaged using a ChemiDoc MP Imaging System (Bio-Rad).

### On-bead digestion and liquid chromatography mass spectrometry

Mass spectrometry-based proteomic experiments were performed as previously described with minor modifications^38^. Briefly, after enrichment and washing, beads were resuspended in 200 μL on-bead digestion buffer (50 mM HEPES pH8.0, 1 μM CaCl_2_, 2% ACN); 10 mM TECP and 40 mM CAA were added and incubated for 30 min at room temperature. The beads were washed with 1 mL on-bead digestion buffer. The beads were resuspended in 100 μL on-bead digestion buffer with 1 μL 0.5 μg LysC (Wako, 125-05061) and incubated at 37°C for 3 h. Then, on-bead digestion buffer with 0.5 μg trypsin (Promega, V5280) was added for digestion at 37°C for 16 h.

The samples were desalted using stage tips before LC-MS/MS analysis. The stage tips were made of C18 material inserted in 200 μL pipette tips. To desalt the peptide samples, C18 material was washed once with 200 μL acetonitrile, once with 200 μL stage tip buffer B (0.1% (vol/vol) formic acid in 50% (vol/vol) acetonitrile/H_2_O), and twice with 100 μL stage tip buffer A (0.1% (vol/vol) formic acid in H_2_O). Peptide samples were loaded on stage tips and washed twice with 100 μL stage tip buffer A. Finally, peptide samples were eluted with 100 μL stage tip buffer C (0.1% (vol/vol) formic acid in 40% (vol/vol) acetonitrile/H_2_O) and 100 μL stage tip buffer B. The solutions were passed through the stage tips by centrifugation at 500 × *g* for 5 min at room temperature. The elution fractions were collected, and the solution was evaporated from peptide samples in a SpeedVac. Finally, 10 μL stage tip buffer A was added to the samples to perform LC-MS/MS analysis.

### LC-MS/MS analysis

All peptides were reconstituted in 0.1% FA (vol/vol) and separated on reversed-phase columns (trapping column: particle size = 3 μm, C18, length = 20 mm (Thermo Fisher Scientific, P/N 164535), analytical column: particle size = 2 μm, C18, length = 150 mm (Thermo Fisher Scientific, P/N 164534)) on an Ultimate™ 3000 RSLCnano system (Thermo Fisher Scientific, San Jose, CA, USA) coupled to Orbitrap Q-Exactive™ HF (Thermo Fisher Scientific). Peptide separation was achieved using a 60-min gradient (buffer A: 0.1% FA in water, buffer B: 0.1% FA in 80% ACN) at a flow rate of 300 mL/min and analyzed by Orbitrap Q-Exactive™ HF in a data-dependent mode. The Orbitrap Q-Exactive™ HF mass spectrometer was operated in positive ion mode with ion transfer tube temperature 275°C. The positive ion spray voltage was 2.1 kV. Full-scan MS spectra (m/z 350-2000) was acquired in the Orbitrap with a resolution of 60,000. HCD fragmentation was performed at normalized collision energy of 28%. The MS2 automatic gain control (AGC) target was set to 5e4 with a maximum injection time (MIT) of 50 ms, and dynamic exclusion was set to 30 s.

### MS data analysis

#### Protein identification and label-free protein quantification

Raw data were processed with MaxQuant (version 1.6.10.43) and its built-in Andromeda search engine for feature extraction, peptide identification, and protein inference. Mouse reference proteome from UniProt Database (UniProtKB/Swiss-Prot and UniProtKB/TrEMBL, version 2020_12) combined with manually annotated contaminant were applied to search the peptides and proteins. The false discovery rate (FDR) values were set to 0.01, and a match-between-runs algorithm was enabled. After searching, the reverse hits, contaminants, and proteins only identified by one site were removed. Filtered results were exported and further visualized using the statistical computer language Python (version 3.8.3), online gene annotation and analysis tool Metascape (version 2021_02), and the complex network visualizing platform Cytoscape (version 3.8.2).

#### Interacting proteins detection of each specific histone mark

First, raw data were analyzed in MaxQuant using the basic principles as described above. Search results were filtered at 0.01 FDR on precursor and protein group level. The Pearson correlation coefficients (PCC) of all replicates were calculated using the function Series.corr() in Python library pandas (version 1.0.5). Two replicate samples with the highest PCC were retained for further analysis.

For each histone mark, the transformed protein intensities fold change (log2-FC) in the pA-APEX2 experiment compared to the no-H_2_O_2_ control experiment and in the pA-APEX2 experiment compared to the IgG control experiment were calculated. For proteins that were not identified in pA-APEX2 experiments but were identified in no-H_2_O_2_ or IgG controls, the log2-FCs were defined as −100. For proteins identified in pA-APEX2 experiments but not in no-H_2_O_2_ or IgG controls, the log2-FCs were defined as 100. Proteins with log2-FC > 1 were ultimately detached from background proteins in two independent measurements, which are considered to be potential interacting proteins of corresponding histone marks.

#### Functional gene set enrichment and interaction network visualization

All proteins identified previously were mapped to mouse Metascape identifiers via the gene names. Functional gene set enrichment for each histone mark was performed using the “Custom Analysis” function in Metascape (version 2021_02). Min overlaps of 3, 0.01 p-value cutoff, and min enrichment value of 1.5 were used. From all enriched proteins in any of the interactions, the top 20 Gene Ontology Cellular Component terms (Gene Ontology Consortium, 2020) that were significantly enriched in at least three of the interactors were selected.

STRING (version 11.0) interaction confidences with a confidence score of 0.4 and FDR stringency of 0.05 were added as links between identified proteins. Proteins in vital GO terms with a positive log2-FC in histone mark interactors compared to no-H_2_O_2_ and IgG were considered to be a visualization foreground. The network of each specific histone mark interactome was imported into Cytoscape (version 3.8.2) and visualized. Cytoscape (version 3.8.2) was used to layout the potential interacting proteins of each histone mark in pA-APEX2 experiments that were members of vital enriched GO terms. Visualization was based on GO term membership.

## Acknowledgements

This work was supported by a grant from the National Key R&D program of China (grant no. 2019YFA0903803), the Major Program of National Natural Science Foundation of China (grant no. 32090031), the General Program of National Natural Science Foundation of China (grant no. 32070610), the Guangdong Province Fund for Distinguished Young Scholars (grant No. 21050001099), the National Natural Science Foundation of China for Young Scholars (grant no. 32000580), Guangdong Provincial Key Laboratory of Synthetic Genomics (grant No. 2019B030301006), Shenzhen Key Laboratory of Synthetic Genomics (grant No. ZDSYS201802061806209), the Mayo Clinic Cancer Center Eagles Cancer Fund (Z.W.), Mayo Clinic Cancer Center Hematologic Malignancies Program (Z.W.), Mayo Clinic division of Hematology (Z.W.) and a grant from the Mayo Clinic Center for Biomedical Discovery (S.M.O).

## Author contributions

H.G. and Z.W. conceived the study. X.L. and Q.W., designed experiments. N.L. provided mass spectrometry technology platform. W.Z. and W.W. designed and conducted mass spectrometry procedures. J.Z. and H.P. processed mass spectrometry data and conducted statistical analyses. All authors interpreted the data. Z.W., X.L. and Q.W. wrote the manuscript, and K.B., S.O. contributed to revision and editing of the manuscript.

