## Supplementary Figures for "Defining proximity proteomics of post-translationally modified proteins by antibody-mediated protein A-APEX2 labeling"

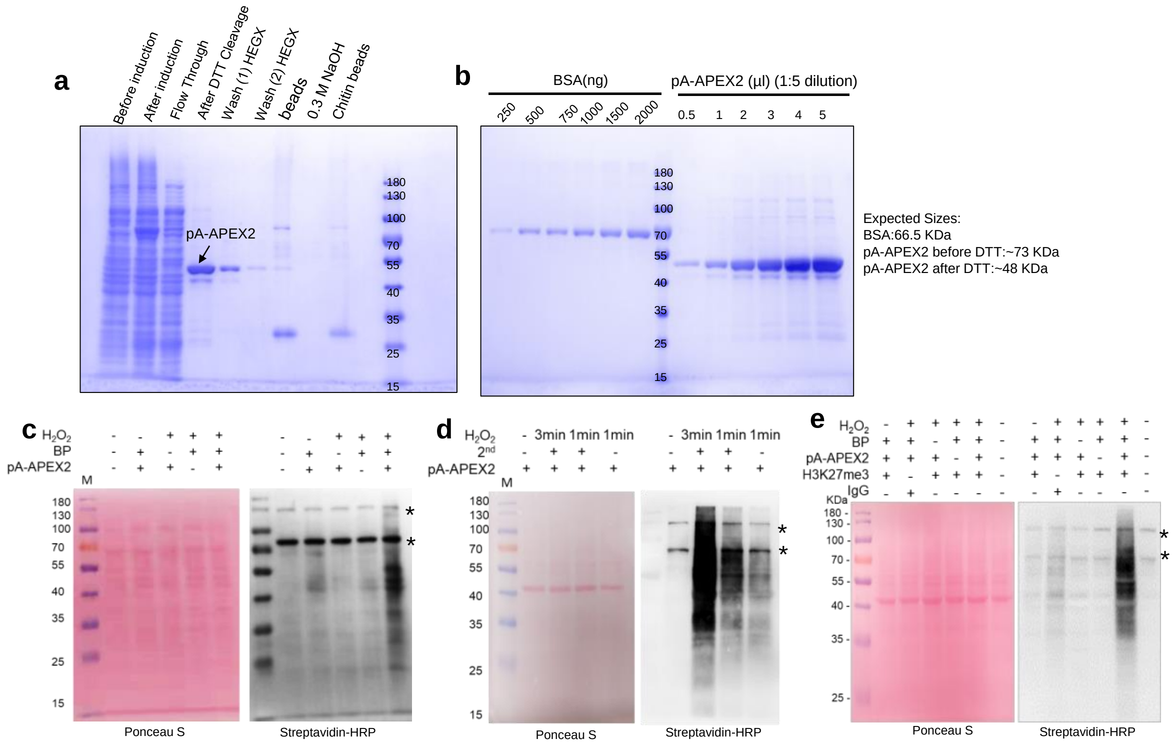


**Supplementary Figure 1. Purified pA-APEX is enzymatically active toward whole-cell lysate.** **a**, Purification process of pA-APEX2. **b**, the concentration of pA-APEX2 was determined by BSA standards, 4μg/μl. **c**, pA-APEX2 labeling *in vitro*. pA-APEX2 and indicated supplements for the peroxidase reaction were incubated with whole-cell lysates as indicated, and the reaction product was analyzed by western blot. **d**, Titration of reaction time and supplements for the pA-APEX2 reaction. **e**, Western blot analysis of H3K27me3-mediated pA-APEX2 proximity labeling protein in living cells by change labeling time or secondary antibody, Whole-cell lysate was analyzed by Ponceau Stain (left) and Western blot (right). "*" denote endogenous biotinylated proteins.

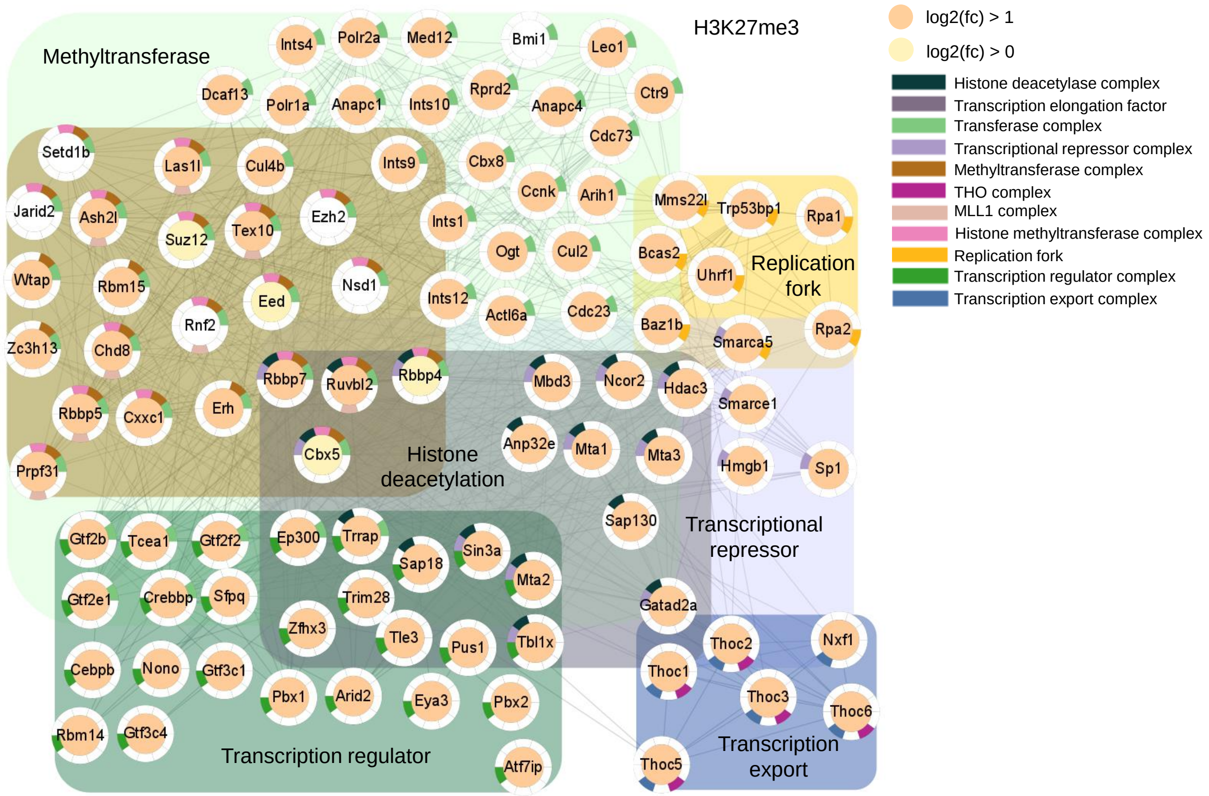


**Supplementary Figure 2**. **The proximal proteome of H3K27me3-interacting proteins identified by AMAPEX.** Network analysis of H3K27me3 interactomes according to the major cellular component GO terms (N proteins = 104, of all 484). Individual proteins are shown as nodes, and interactions are shown as edges. The interactions were retrieved from the STRING database with interaction score > 0.4.

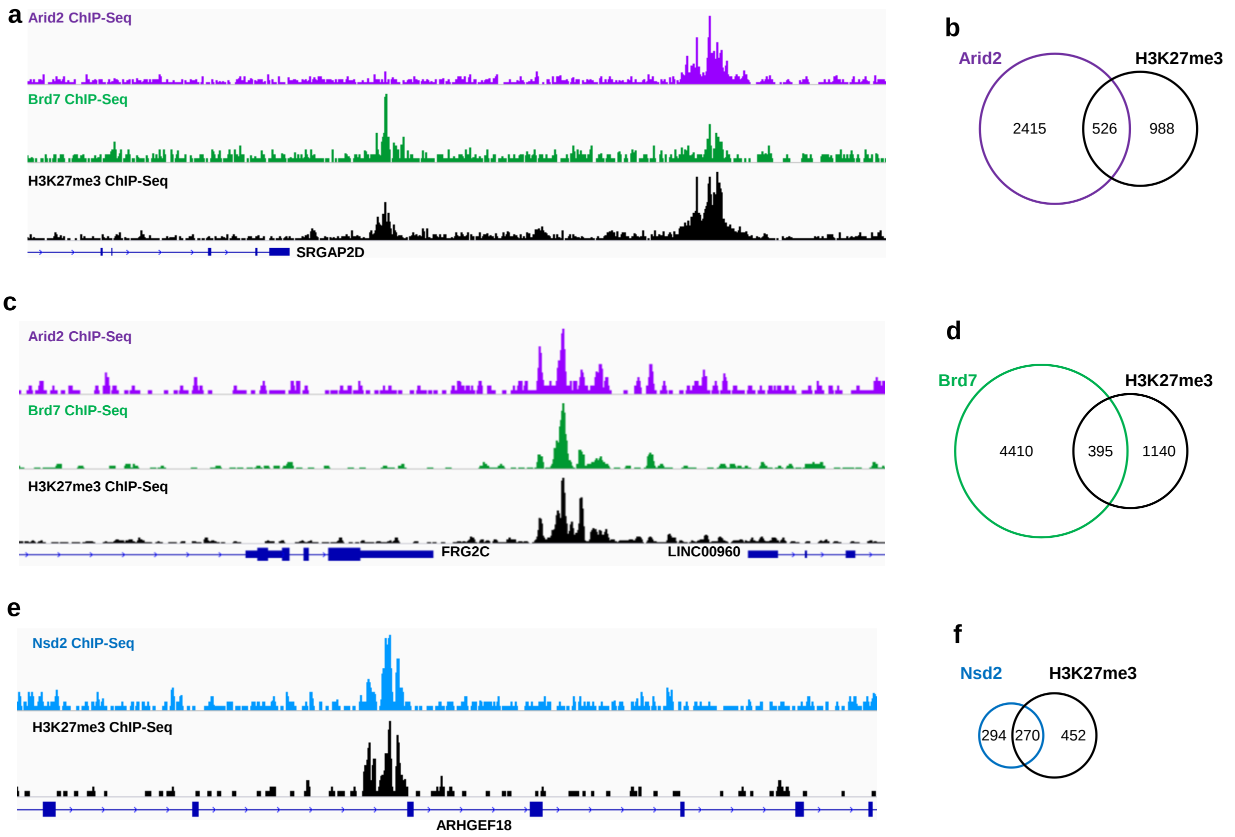


**Supplementary Figure 3. The interactions between H3K27me3 and Arid2, Brd7, or Nsd2 are verified by ChIP-seq**. **a, c, e**, Representative co-localization of ChIP-seq peaks between H3K27me3 and Arid2, Brd7 or Nsd2. **b, d, f**, Venn diagrams showing the number of H3K27me3 peaks overlapping with Arid2, Brd7, or Nsd2 peaks. The ChIP-seq datasets for Arid2, Brd7, and H3K27me3 are from MCF-7 cells. The ChIP-seq dataset for Nsd2 and H3K27me3 are from K-562 cells. All of the bigwig files and peak files were download from https://chip-atlas.org/.

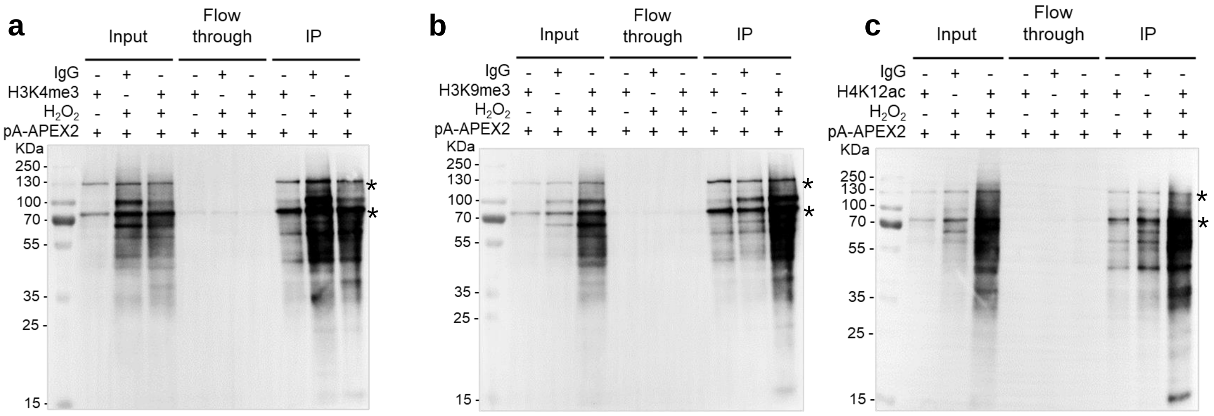


**Supplementary Figure 4. Antibody-mediated specific biotin labeling by pA-APEX2.**

MEF cells were briefly crosslinked with 0.1% formaldehyde before the antibody-directed pA-APEX2 biotinylation reaction. Whole cell lysates were extracted, and biotinylated proteins were purified using streptavidin beads. Whole-cell lysates (input), flow throughs, and immunoprecipitated (IP) proteins were analyzed by western blot. **a**, H3K4me3, **b**, H3K9me3, **c**, H4K12ac. “*” denote endogenous biotinylated proteins.

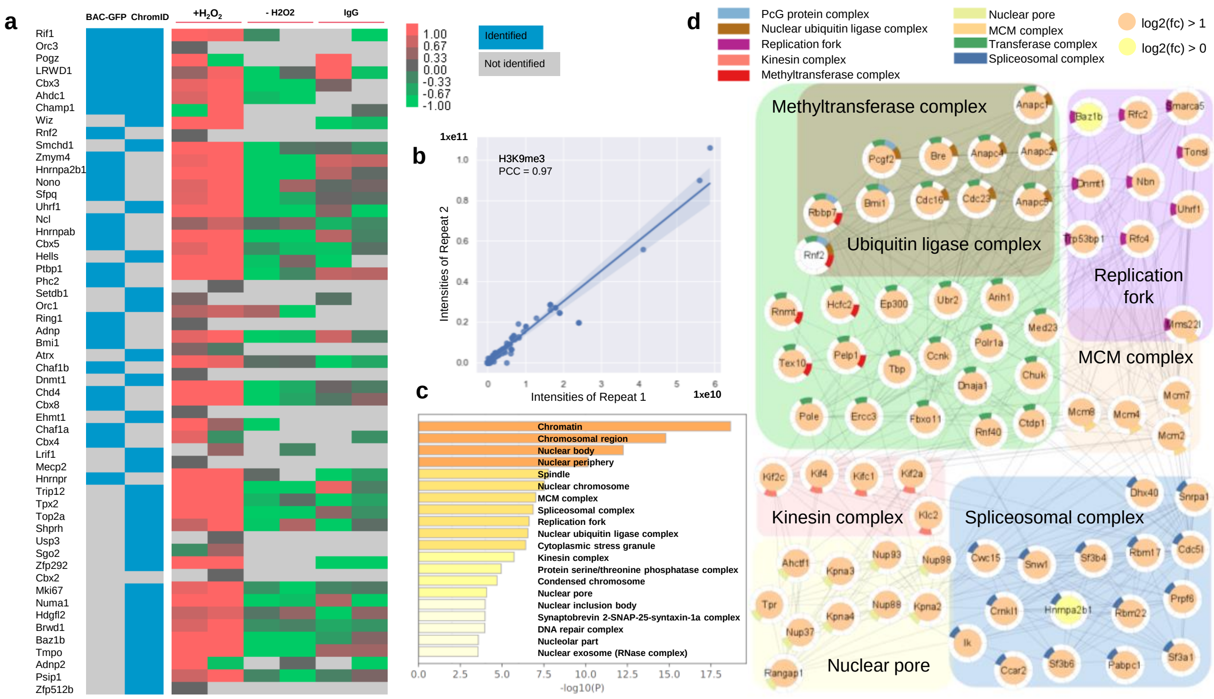


**Supplementary Figure 5. The proximal proteome of H3K9me3 identified by AMAPEX.**

**a**, H3K9me3-interacting proteins identified by AMAPEX. Heatmap showing the enrichment as log2-fold intensity change of interacting proteins relative to the controls as indicated. Data are shown as Z scores. Blocks in blue represent the enrichment of proteins identified by ChromID and BAC-GFP in previous publications. **b,** The reproducibility of two biological replicates of pA-APEX2 experiments in the identification of H3K9me3-interacting proteins was calculated by Pearson correlation coefficient (PCC). **c,** The top 20 enriched cellular component GO terms for the H3K9me3 proximal proteins. Bar plots represent the -log10 (p value) of the enriched terms. **d**, Network analysis of H3K9me3 interactomes according to the major cellular component GO terms (N proteins = 83, of all 486). Individual proteins are shown as nodes, and interactions are shown as edges. The interactions were retrieved from the STRING database with interaction score > 0.4. Proteins were selected based on min 1.0 log2-FC in two pA-APEX2 experiments. Proteins detected in pA-APEX2 experiments with log2-FC > 1 are shown as orange nodes, proteins with log2-FC > 0 and detected in ChromID or BAC-GFP are shown as light-yellow nodes, and proteins detected in ChromID or BAC-GFP but not in pA-APEX2 experiments are shown as white nodes.

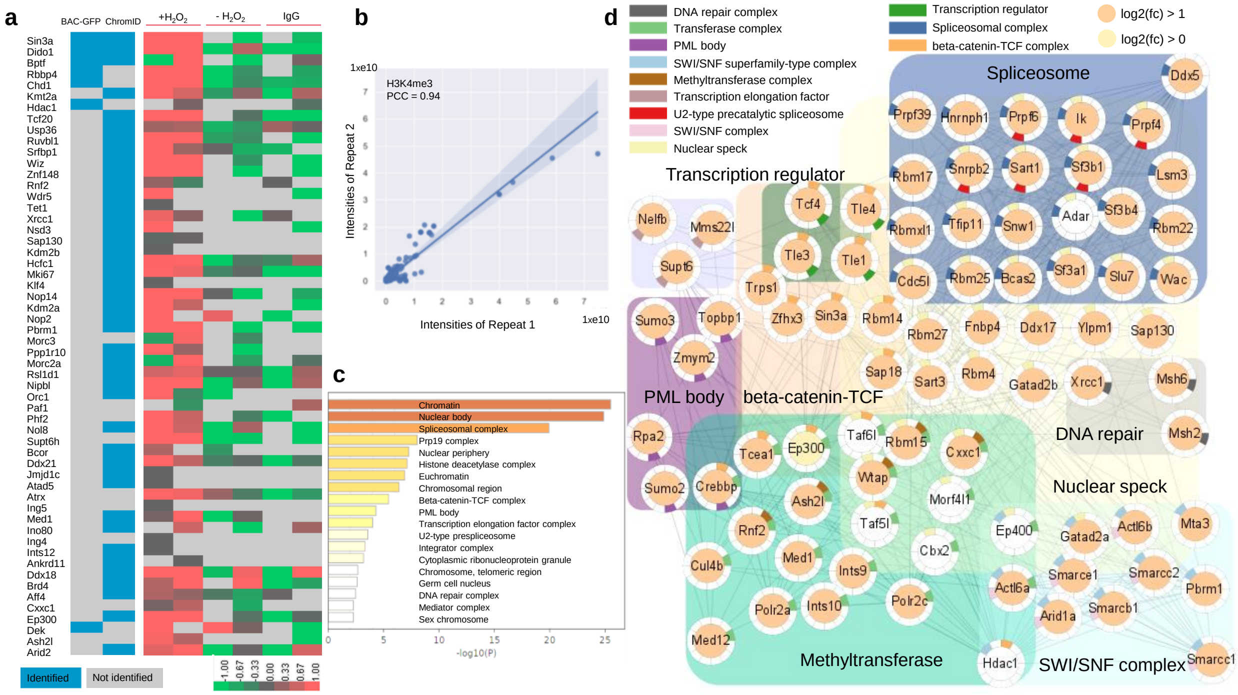


**Supplementary Figure 6. The proximal proteome of H3K4me3 identified by AMAPEX.**

**a,** H3K4me3-interacting proteins identified by AMAPEX. Heatmap showing the enrichment as log2-fold intensity change of interacting proteins relative to the controls as indicated. Data are shown as Z scores. Blocks in blue represent the enrichment of proteins identified by ChromID and BAC-GFP in previous publications. **b**, The reproducibility of two biological replicates of pA-APEX2 experiments identifying H3K4me3-interacting proteins. The Pearson correlation coefficient (PCC) between two replicates was calculated. **c,** The top 20 enriched cellular component GO terms for the H3K4me3-proximal proteins. Bar plots represent the -log10(p value) of the enriched terms. **d**, Network analysis of H3K4me3 interactomes according to the major cellular component GO terms (N proteins = 91, of all 193). Individual proteins are shown as nodes, and interactions are shown as edges. The interactions were retrieved from the STRING database with interaction score > 0.4. Proteins were selected based on min 1.0 log2-FC in two pA-APEX2 experiments. Proteins detected in pA-APEX2 experiments with log2-FC > 1 are shown as orange nodes, proteins with log2-FC > 0 and detected in ChromID or BAC-GFP are shown as light-yellow nodes, and proteins detected in ChromID or BAC-GFP but not in pA-APEX2 experiments are shown as white nodes. H3K4me3-interacting proteins identified by pA-APEX2 experiments indicate that the interactome is enriched for spliceosome, transcription regulation, and methyltransferase complexes.

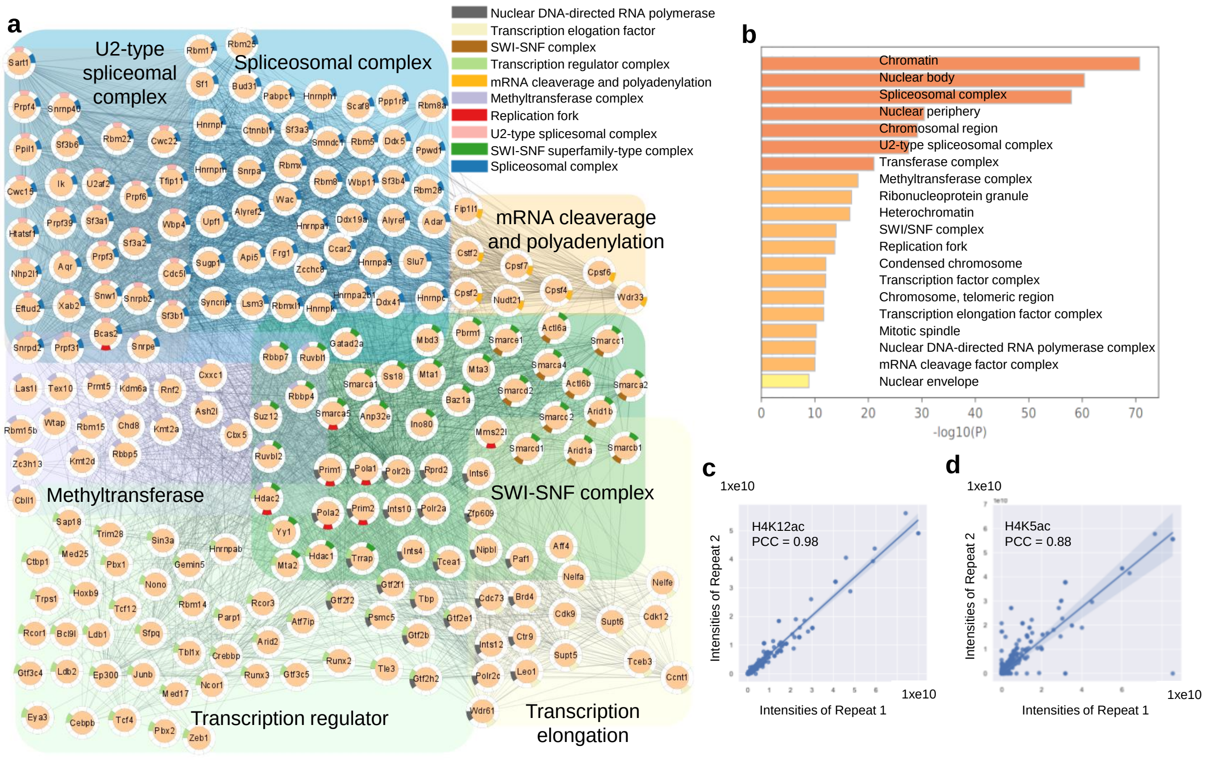


**Supplementary Figure 7. The proximal proteome of H4K12ac identified by AMAPEX.**

**a**, Network analysis of H4K12ac interactomes according to the major cellular component GO terms (N proteins = 215, of all 1,019). Individual proteins are shown as nodes, and interactions are shown as edges. The interactions were retrieved from the STRING database with interaction score > 0.4. Proteins were selected based on min 1.0 log2-FC in two pA-APEX2 experiments. **b**, The top 20 cellular component GO terms for the H4K5ac-proximal proteins. Bar plots represent the -log10(p value) of the enriched terms. **c**, d, The reproducibility of two biological replicates of pA-APEX2 experiments identifying H4K12ac/H4K5ac-interacting proteins. The Pearson correlation coefficient (PCC) between two replicates was calculated.

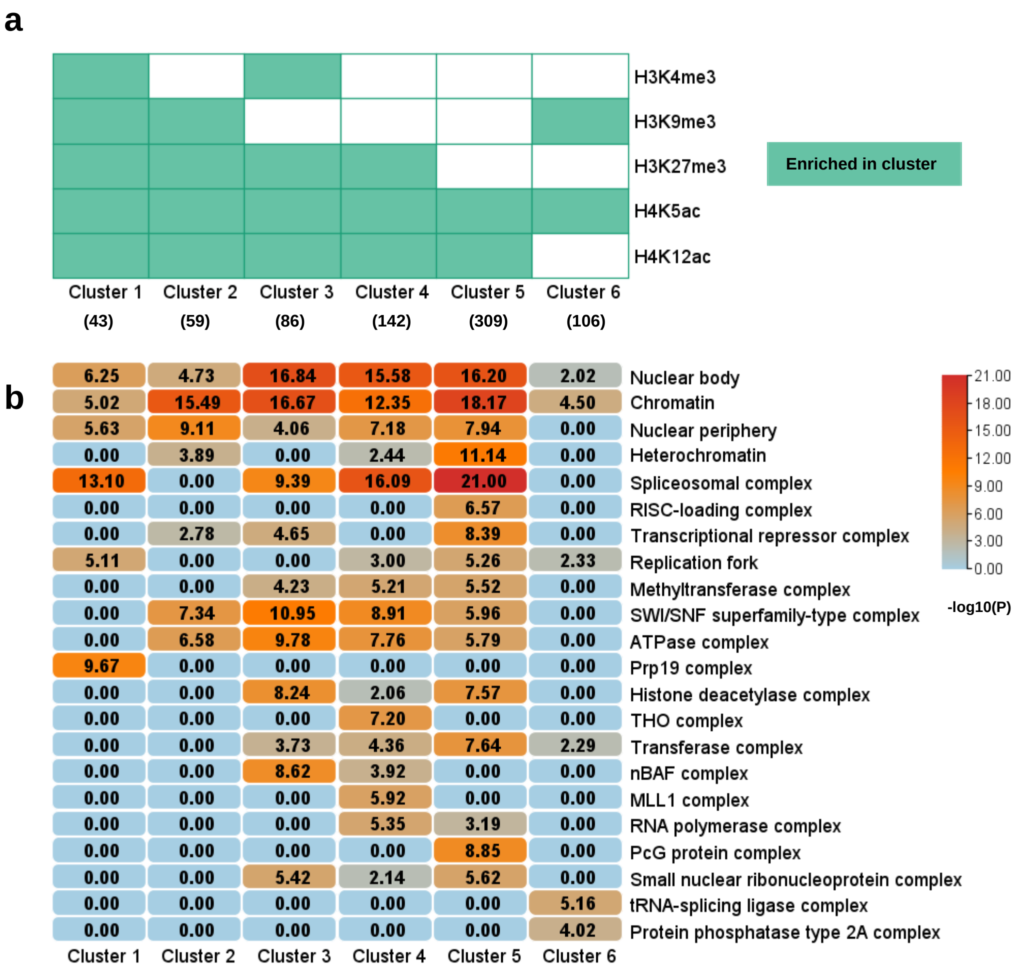


**Supplementary Figure 8. Functional clusters of interacting proteins of five histone marks identified by AMAPEX. a**, Proximal proteins of five histone marks were selected based on their log2-FC, and overlapping proteins between each histone mark were clustered using Gene Cluster 3.0. Blocks in green represent interacting proteins of specific histone marks that are enriched in the corresponding cluster. Numbers of proteins are shown in parentheses. **b**, Functional enrichment analysis based on major cellular component GO terms identified in six clusters. Metascape was used for the enrichment analysis (min overlap = 3, p-value = 0.01, and min enrichment value = 1.5). The -log10(p value) was adopted to infer the enrichment score of each cluster.
